# i-stLearn: An interactive platform for spatial transcriptomics analysis

**DOI:** 10.1101/2023.03.27.534291

**Authors:** Duy Pham, Brad Balderson, Quan H. Nguyen

## Abstract

**Summary:** Emerging spatial transcriptomics technologies (e.g. Visium, Slideseq, or MERFISH) have made it possible to keep the spatial information while profiling gene expression of every cell/spatial-spot. Integrating expression values, spatial coordinates, and imaging data type promises to bring more biological insights but is still technically challenging. A user-friendly software tool to enable interactive analysis of spatial transcriptomic data by the broader community is lacking. We present i-stLearn, an all-in-on web application with an analysis pipeline and interactive visualization for studying spatial heterogeneity using spatial transcriptomics data. i-stLearn can be used to gain biological insights from tissue through key analysis types cell-cell interaction analysis, clustering, and trajectory inference. Using functions, users can interactively segment the tissue and identify cellular state transition or cellular communications in a heterogeneous biological sample.

**Availability:** i-stLearn is freely available at https://github.com/BiomedicalMachineLearning/stlearn_interactive as local web application and we also provide a demo online web service available at https://i-stlearn-demo.web.app.

**Supplementary information:** Supplementary data are available at *Bioinformatics* online.

## 1 Introduction

Spatial transcriptomics has been developed to preserve spatial dimensional, while generating single-cell resolution or spot-level transcriptional profiles [1, 2]. Technologies like Visium also add tissue morphology images. The rapid development of spatial technologies also leads to an urgent need for analysis tools available to the broad research community to utilise present, and future spatial data sets [3].

We developed a comprehensive spatial analysis pipeline, implemented in an interactive software application, named as i-stLearn. This software provides a thorough downstream analysis, including pre-processing (exploratory analyses and normalisation), clustering, spatio-temporal trajectory/transitioning analysis and spatial cell-to-cell interaction analysis. For all steps, i-stlearn provides interactive options, fully enabled with histological tissue visualisation functionality. i-stLearn allows users to select specific tissue regions for deeper analysis, an important feature that is lacking in many spatial transcriptomics analysis tools. To ensure reproducibility,log files are generated.

The source code is freely available at: https://github.com/BiomedicalMachineLearning/stlearn interactive and the demo website is at: https://i-stlearn-demo.web.app.

## 2 i-stLearn functions and interactive visualizations

### 2.1 Preparing data input

i-stLearn provided two options for users to upload their data. Due to the popularity of Visium platform data (10X genomics), users can upload the Space Ranger output folder directly to i-stLearn web application. For other platforms or Visium processed data, we support users to upload h5ad files which store the common AnnData object [4] and this object should be created with stLearn [2] (Figure 1A).

**Fig. 1.**
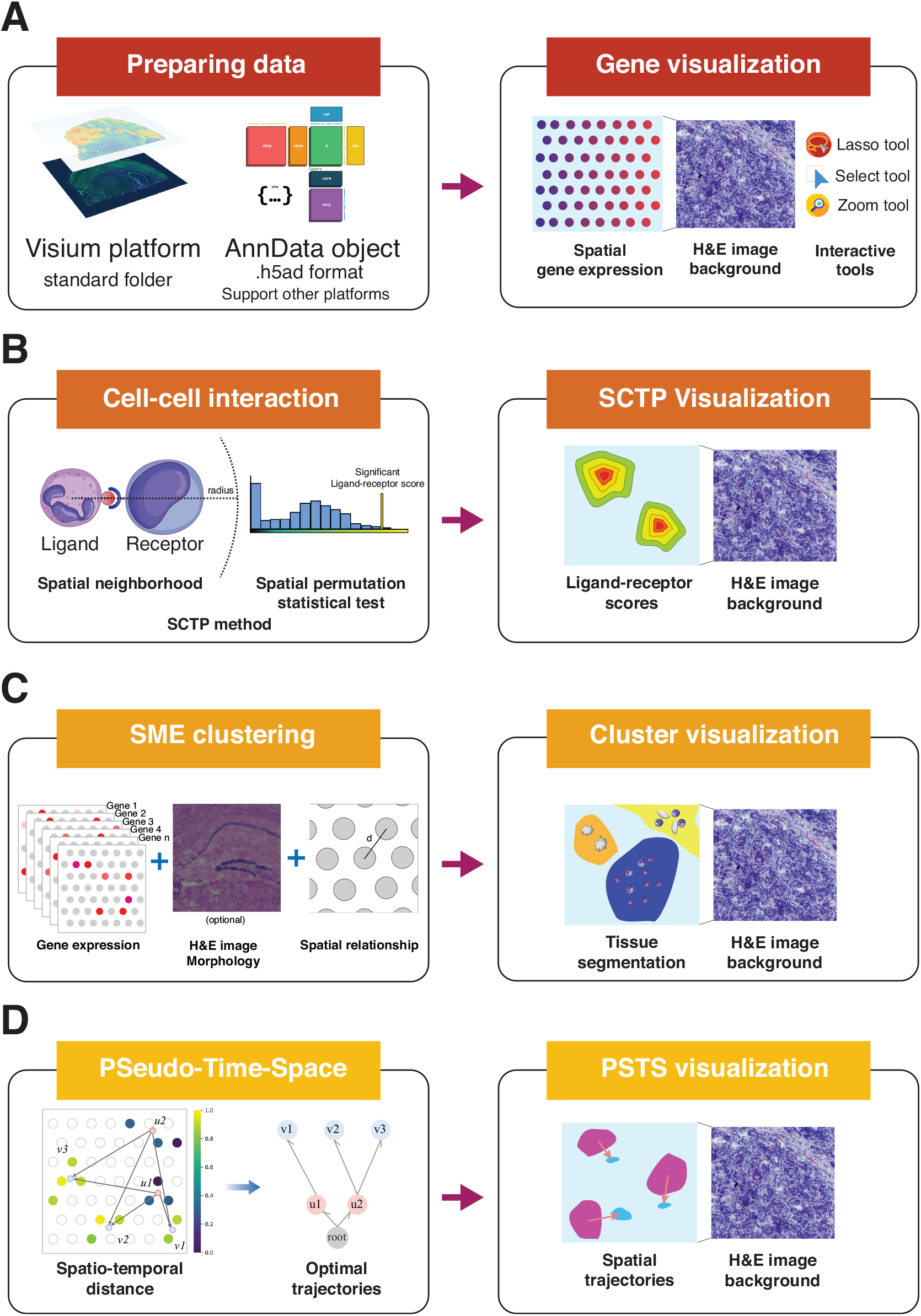
i-stlearn an interactive web application for spatial transcriptomics analysis. **A**. i-stLearn takes input as the Visium CellRanger output (for Visium data) or AnnData object (Visium or other types of data). The application provides interactive plotting functions to visualize the gene expression with the H&E image background and precise spatial coordinates. i-stLearn also includes interactive selection/visualisation functions like Lasso, select, or zoom tools. **B**. The cell-cell interaction analysis can be run with the SCTP method that incorporates spatial neighborhood and a permutation statistical test. Users can visualize the ligand-receptor interaction scores within the tissue image. **C**. For clustering, i-stLearn provides a SME clustering option, which combines gene expression and spatial distance information (optionally with morphological information) to improve the detection sensitivity. Users can see the clustering results overlayed on tissue segmentation result at the background. **D**. i-stLearn also implements PSTS algorithm for trajectory inference, which uses spatio-temporal distance to optimize the trajectories and users can view the spatial trajectory results on the tissue section or across the spatial coordinates.

### 2.2 Gene expression visualization

Gene expression plots and all other interactive plots in i-stLearn are made using Bokeh (Figure 1A). i-stLearn includes powerful tools from Bokeh like Lasso select, box select, and wheel zoom, allowing users to customise and save plots. We also added multiple options to work directly with the data and enable on-the-fly updating of the data for plotting like gene selection, spot/cell opacity, size and colour map.

### 2.3 Cell-cell interaction analysis

Users can predict the cell-cell interaction with the ligand-receptor pairs scoring by using our provided function (Figure 1B). We built it based on the spatially-constrained two-level permutation (SCTP) method [2] from the stLearn toolkit.

### 2.4 SCTP visualization

i-stLearn allows users to observe the ligand-receptor interaction scores across the tissue, as calculated by SCTP method (Figure 1B). The score is proportional to the level of predicted interaction. For visualization, we use the same function as the gene expression interactive plot.

### 2.5 Clustering using data from Spatial Morphology Expression correction/imputation - SME clustering

In i-stLearn, we set an option to allow users to optionally perform SME correction/imputation step [2] (Figure 1C). This correction/imputation makes use of three types of information, imaging of tissue morphology, spatial distance and gene expression. Post SME, users can use multiple methods in i-stLearn to perform clustering.

### 2.6 Cluster visualization

i-stLearn supports the views of any set of clusters users are interested in displaying (Figure 1C). To assess clustering results, we also provide a function to display dot matrix plots for visualizing the differential expression analysis result for each cluster.

### 2.7 Pseudo-time-space analysis

To reconstruct the spatial trajectories on a tissue section, i-stLearn takes the clustering result as the initial step for pseudo-time-space (PSTS) method [2] (Figure 1D). For the parameters setting, users can select the root cluster, and available paths from the spatial PAGA graph [2] (Partition-based graph abstraction) as a base connection between clusters.

### 2.8 Spatial trajectories visualization

i-stLearn offers an innovative visualization of the spatial trajectory results using the cluster plot function (Figure 1D). In the interactive plot, spatial trajectories are shown as arrows that connect spots/cells from root sub-clusters to target sub-clusters.

## 3 Conclusion

i-stLearn is an all-in-one and user-friendly spatial transcriptomics data analysis pipeline and interactive visualization. We trust that i-stLearn will contribute to enabling the broad research community to utilize spatial transcriptomics data.

## References

[1] Lewis, S.M., Asselin-Labat, M.-L., Nguyen, Q., Berthelet, J., Tan, X., Wimmer, V.C., Merino, D., Rogers, K.L., Naik, S.H.: Spatial omics and multiplexed imaging to explore cancer biology. Nature methods 18(9), 997–1012 (2021)

[2] Pham, D., Tan, X., Xu, J., Grice, L.F., Lam, P.Y., Raghubar, A., Vukovic, J., Ruitenberg, M.J., Nguyen, Q.: stlearn: integrating spatial location, tissue morphology and gene expression to find cell types, cell-cell interactions and spatial trajectories within undissociated tissues. BioRxiv (2020)

[3] Moses, L., Pachter, L.: Museum of spatial transcriptomics. Nature Methods 19(5), 534–546 (2022)

[4] Virshup, I., Rybakov, S., Theis, F.J., Angerer, P., Wolf, F.A.: anndata: Annotated data. bioRxiv (2021)5

