## Supplementary document 1 for "i-stLearn: An interactive platform for spatial transcriptomics analysis"

### 1 Urgent need of spatial transcriptomics analysis tools

Traditional RNA sequencing and single-cell RNA sequencing technology have become the methods of choice for researchers to study the composition of cell types within each tissue throughout the past decade[1,2]. However, such sequencing technologies discard spatial information[3,4]. To address this critical constraint,

So far, just a few toolkits have been created specifically for spatial transcriptomics data, most of which are still based on traditional single-cell analysis methods without making use of spatial or imaging information[5,6]. Additionally, user-friendly spatial software toolkits are still lacking for most biologists who are providing expert-domain understandings but are not familiar with command-line analysis.

### 2 i-stLearn architecture

We designed i-stLearn to serve the broad research community by making the analysis easy from installation to operation. i-stlearn web application has two parts, backend (computation) and frontend (web interface). For the backend, the main analysis pipeline is based on the stLearn[7] and Scanpy[8] toolkits, which are designed for the spatial and scRNAseq data analysis respectively. All functions are managed under the Flask web framework[9]. For the frontend, we used Bokeh[10] for interactive visualization of spatial transcriptomics data. This capability is especially useful for a histopathological workflow. Users will be able to upload raw data for processing with the i-stlearn pipeline or processed data for visualization and further spatially-aware analysis.

#### 2.1 Backend

The main web framework that we used is Flask[9] because it islightweight, yet provides us with all essential tools and features to create web applications. The Flask app file creates and controls all the computational Application Programming Interface (API) and interactive plotting API.

For the computational backend, we defined several APIs such as: "/upload" - uploading data (support both Visium folder and H5AD anndata[8] object file); "/preprocessing" - filter gene and cell/spot, normalize, log transform and scale data; "/clustering" - perform clustering with an option for imputation using the integrative modelling of Spatial, Morphological, and Gene expression data (SME)[7]; "/lr" - ligand receptor interaction analysis with a Spatially Constrained Two Level Permutation test (SCTP)[7]; "/cci" cell-cell interaction analysis with scanning neighborhood method; "/psts" - pseudo-time-space method[7] for spatial trajectories inference; and "/dea" - differential expression analysis[8]. Besides, we used WTForms which is a library for flexible form validation and rendering to manage all the parameter settings of computational API to enable reproducibility for analysis workflow by users.

One important part of the analysis backend are the three Python libraries, including anndata, scanpy[8] and stLearn[7]. For broad applicability, we used the main object as AnnData object from anndata library. All the spatial transcriptomics data is stored in that object, and all provided functions interact with it. On top of that, we used scanpy and stLearn for data processing parts. For example, *scanpy.read_visium* is used for reading Visium folder or $stlearn.tl.cci.run$ is used for running the ligand-receptor analysis.

For the interactive plotting backend, we used Bokeh[10] to generate and manage the parameter setting. Bokeh library provided users with control options using text box, slice and checkbox selection for the form to update the interactive plots. This setup allows users to input relevant information to quickly and interactively change the plot, for example, gene selection input as a text box which can help to select the gene to visualize its expression values in the spatial context.

For supporting functions, differential expression analysis (DEA) is a crucial downstream step after we annotated a dataset[11]. i-stLearn also provides a function to perform the DEA with different methods like t-test and Wilcoxon test.

#### 2.2 Frontend

Javascript in the jinja template[12] of Flask is the programming language we used for the frontend component. However, the main interactive plots are built based on the Bokeh library.

The Bokeh plots include several tools that help to interactively work with the plots, such as lasso and box selection, zoom in, zoom out, hover tool, etc. These tools do not directly control the data so the performance is fast. Additionally, we support different colormap schemes, dot sizes, arrow types, etc to add more flexibility for users to analyze the spatial transcriptomics data. Some static plots are also utilized like violin plots and dot-matrix plots for differential expression analysis results.

With the interactive plots, this is especially useful for spatial transcriptomics analysis where users can zoom in to a specific region of interest within the H&E image (e.g. tumour infiltrating cancer region) and check the gene expression in that region.

Notably, to allow for reproducibility, i-stLearn has saving options of plots and data, together with producing running log files that report options selected by users throughout the pipeline. In addition, if clustering information is available in the AnnData object, users can use i-stLearn to produce a violin plot to show the distribution of genes in each cluster, a convenient function for assessing the clusters interactively and quantitatively.

### 3 Breast cancer case study

Non-invasive ductal carcinoma *in situ* (DCIS) is the earliest-detectable form of breast cancer in mammograms[13,14]. Although not all detected DCIS would progress to invasive ductal carcinoma (IDC), consistent predictors for future invasiveness remain elusive, thereby complicating clinical management in terms of treatment[15]. To answer this question, we will apply the computational analysis pipeline that we constructed in i-stLearn (Figure S1).

#### 3.1 Reading data

The breast cancer data is the DCIS sample from an open-source dataset of 10X genomics. The format is the output as the Visium folder of Spaceranger software. We read spatial transcriptomics data input by using the uploading API of i-stLearn. The uploaded API will store all the data in an AnnData object.

#### 3.2 Preprocessing

Here, we performed quality control by running the genes filter to keep genes detected in at least 3 cells/spots. Next, the variance stabilization using log1p transformation and data scaling steps were applied to perform the standard preprocessing for the spatial transcriptomics dataset.

#### 3.3 Ligand-receptor interaction analysis

In this step, we perform the cell communication analysis using the SCTP algorithm from stLearn. By default, only the ligand-receptor (LR) pairs that are co-expressed between neighbour cells in more than 20 locations across the dataset were kept for the analysis. We generated 10,000 pairs (default) to establish the background signal to perform the permutation test for LR pairs with signal significantly higher than the background. LR scores of significant spots can be visualised in the "LR plot" tab (Figure S1E).

The SCTP method truly uses spatial information to reduce false positive detection of cell-cell interactions. Both human and mouse ligand-receptors databases for screening are available for choosing in i-stLearn. For the parameter setting, the higher number of random gene pairs (we recommend 10,000 pairs) for the permutations test, the longer the computation time, but in return, the robustness of the test is improved.

#### 3.4 Clustering

We performed clustering with SME imputation method from stLearn. SME can recover signal loss due to the high dropout rate in spatial sequencing. SME can also predict gene expression in tissue regions not covered by spatial spots. Briefly, PCA results from dimensonality reduction from gene expression matrix were integrated with the morphological features extracted from the H&E image by SME. We then applied Louvain[16] clustering using the SME-adjusted PCA space to segment spots in the breast cancer tissue. Depending on the dataset, we can select parameter settings to fit the biological cell types or tissue architecture. As the results, 12 clusters were defined to represent the breast tissue architecture (Figure S1A). We run the differential expression analysis (DEA) module and identified the gene markers for each of the cluster. i-stLearn allows us to plot both clustering results and DEA results in the "Cluster plot" tab. Moreover, a violin plot in the "Gene plot" tab will show the distribution of the selected gene across all clusters. We will focus on two important clusters that related to our research question: DCIS (6 - pink) (Figure S1B) and IDC - immune infiltration (8 - blue sky) (Figure S1C). We observed that KRT5, KRT14 (We show only KRT14 as an example of visualization in the Figure S1D) and KRT17 are significant in the DCIS cluster; KRT37 and MYO3B are significant in the IDC - immune infiltration cluster. Also, i-stLearn allows the plotting of the number of top genes and min log fold-change. For other interactive functions, this clustering plot is similar to gene expression and ligand-receptor plots.

#### 3.5 Spatial trajectories inference

We applied the pseudo-time-space (PSTS) algorithm in the "PSTS analysis" tab to build the spatial trajectories from DCIS to IDC cluster. First, i-stLearn allows us to choose the root cluster as DCIS and automatically screens all the possible paths that are connected between clusters, where the connections are defined based on gene expression with spatial PAGA graph. Spatial PAGA is based upon PAGA[17], but combines spatial information and gene expression to generate a topology-preserving map of spots/cells. The results will be multiple sub-graphs that represent multiple spatial trajectories of spatial sub-clusters. Next, we selected the path to be the trajectory to calculate pseudotime values. Finally, we reconstructed the spatial trajectories based on both pseudotime values and spatial relationship between clusters (by spatial sub-clusters). As the results, we predicted three spatial trajectories from DCIS to IDC and displayed it in the "Cluster plot" tab (Figure S1F).

#### 4 Slide-seq case study

In this section, we show an example of Slide-seq data to demonstrate that i-stLearn can support different types of spatial transcriptomics platforms.

##### 4.1 Reading data

For Slide-seq[18] or other spatial transcriptomics platforms, we support the AnnData object in the format of an H5AD file. To prepare this file, we should use the stLearn library to create the AnnData data object by count matrix and the spatial data and then export to the H5AD format file to be ready to upload to i-stLearn.

##### 4.2 Analysis

Here, we only demonstrate the basic pipeline to show the use of i-stLearn in a non-visium platform. In the pipeline, we used all the preprocessing steps as filtering cell/spots/genes, normalization, log1p transformation and data scaling. After that we perform the Leiden clustering[16] (Figure S2A) and DEA (Figure S2B) to find the markers for each cluster result (Figure S2C, D, E).

#### 5 Conclusion

In summary, i-stLearn is an all-in-one and user-friendly spatial transcriptomics data analysis pipeline and interactive visualization. Users can use the software to perform preprocessing, cell-cell interaction analysis, clustering, differential expression analysis, and pseudo-time-space analysis making use of spatial information. In addition, various interactive visualization in i-stLearn will be helpful in understanding the analysis results. Users can run the interactive analysis via installing to the local computer and we also implemented a demo as the online service in https://i-stlearn-demo.web.app to preview all functions and the result of the analysis pipeline in a breast cancer dataset. Detailed documentation is provided. We trust that i-stLearn will contribute to enabling the broad research community to utilize spatial transcriptomics data.

We also describe two case studies as above and we hope users will find them useful for using i-stLearn. In additional, we provided a tutorial from Figure S3 to Figure S12.

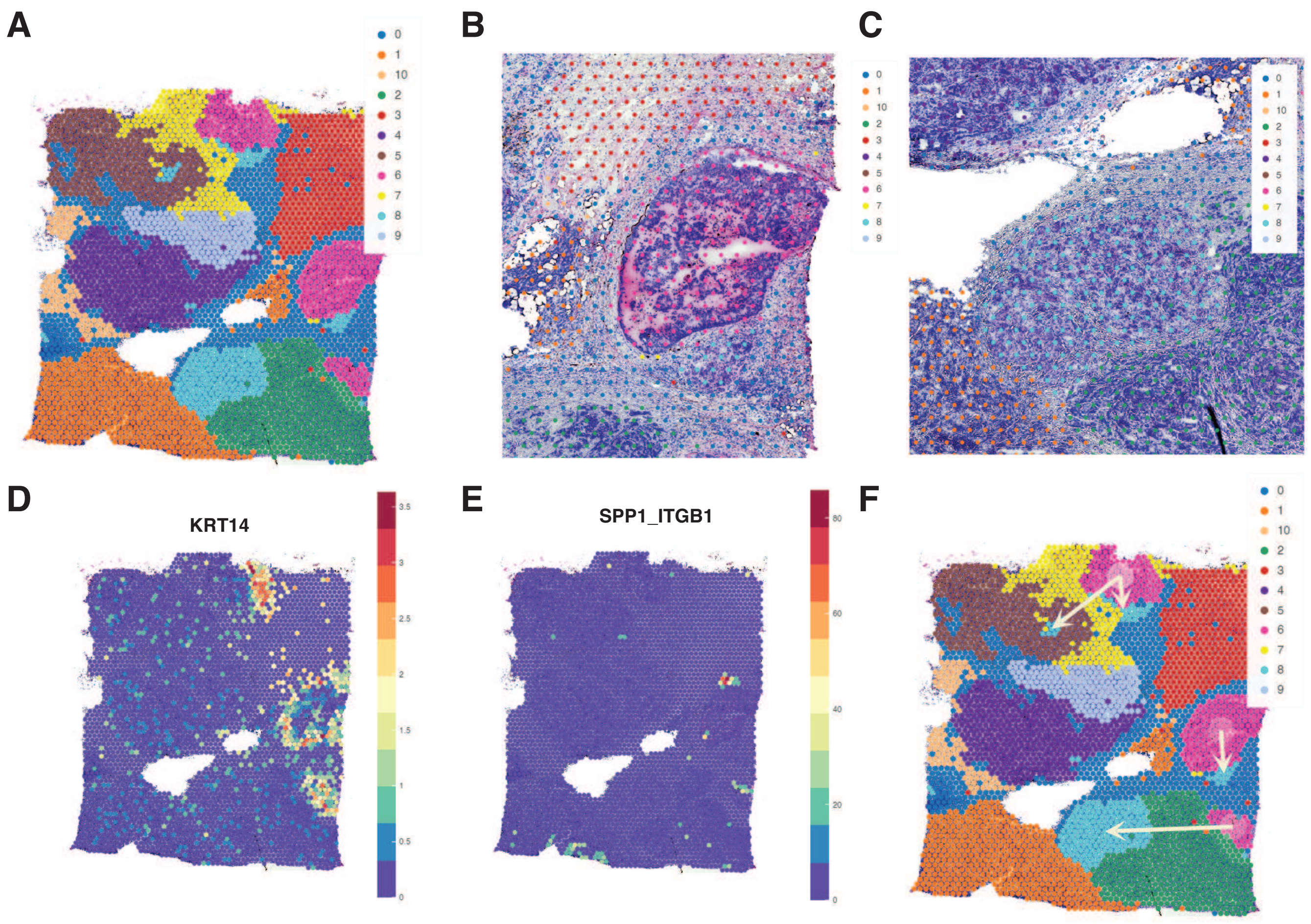


**Supplementary Figure 1. Breast cancer analysis with Visium - spatial transcriptomics data**. **A**, Clustering results reflect the tissue architecture of breast cancer. 11 clusters are found. **B**, Zoom in selection for the regions of ductal carcinoma in situ (DCIS). **C**, Zoom in selection for the regions of invasive ductal carchinoma (IDC). **D** Visualisation of the expression of the KRT14, a biomarker of DCIS region. We can observe this gene in the peripheral edge of the DCIS region. **E**, We predicted the location and the score for the coexpression of the SPP1-ITGB1 ligand-receptor pair. The distribution is at the edge section of the DCIS region. **F**, The arrows display the reconstructed spatial trajectories predicted by PSTS. These trajectories are connected between DCIS and IDC by 3 different clades.

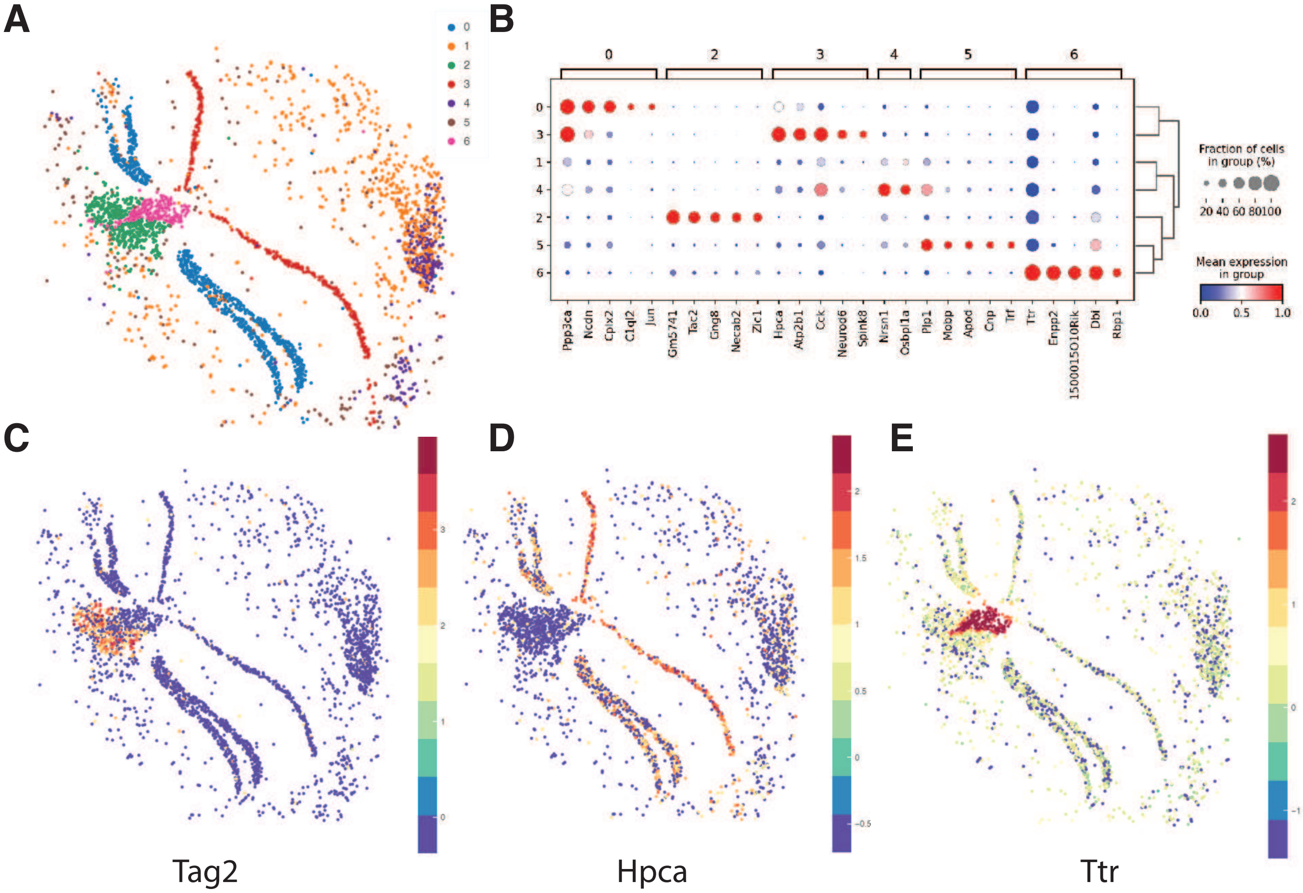


**Supplementary Figure 2.** **Mouse brain analysis with Slide-seq spatial transcriptomics data**. **A**, Clustering results of the mouse brain. **B**, Differential expression analysis results are shown as a dot matrix. Marker expression is shown for each cluster. **C**, Expression of Tag2, a marker of the Habenula part in the brain. **D**, Expression of Hpca, a marker for the CA1 layer in the hippocampus. **E**, Expression of Ttr, a marker for the choroid region in the mouse brain.

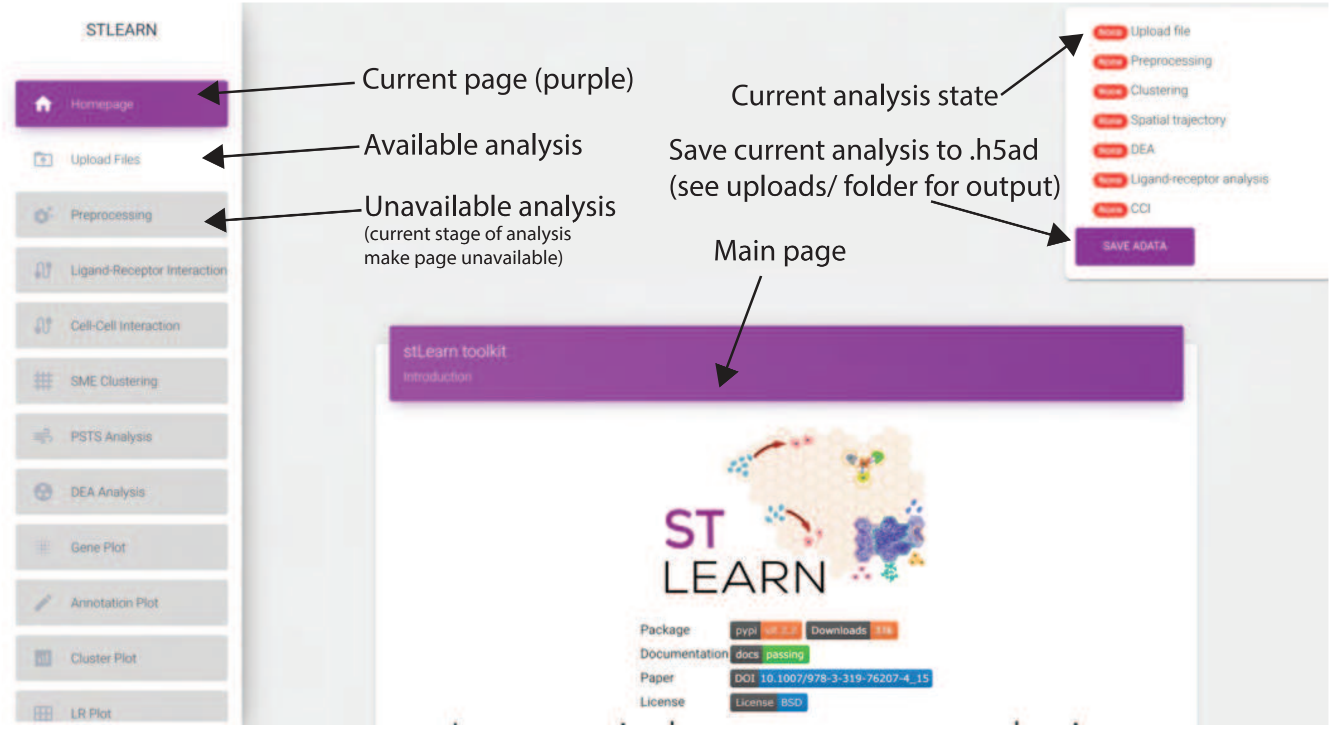


**Supplementary Figure 3.** **A general view of i-stLearn interface and the useful display of the status for data analysis progress**.

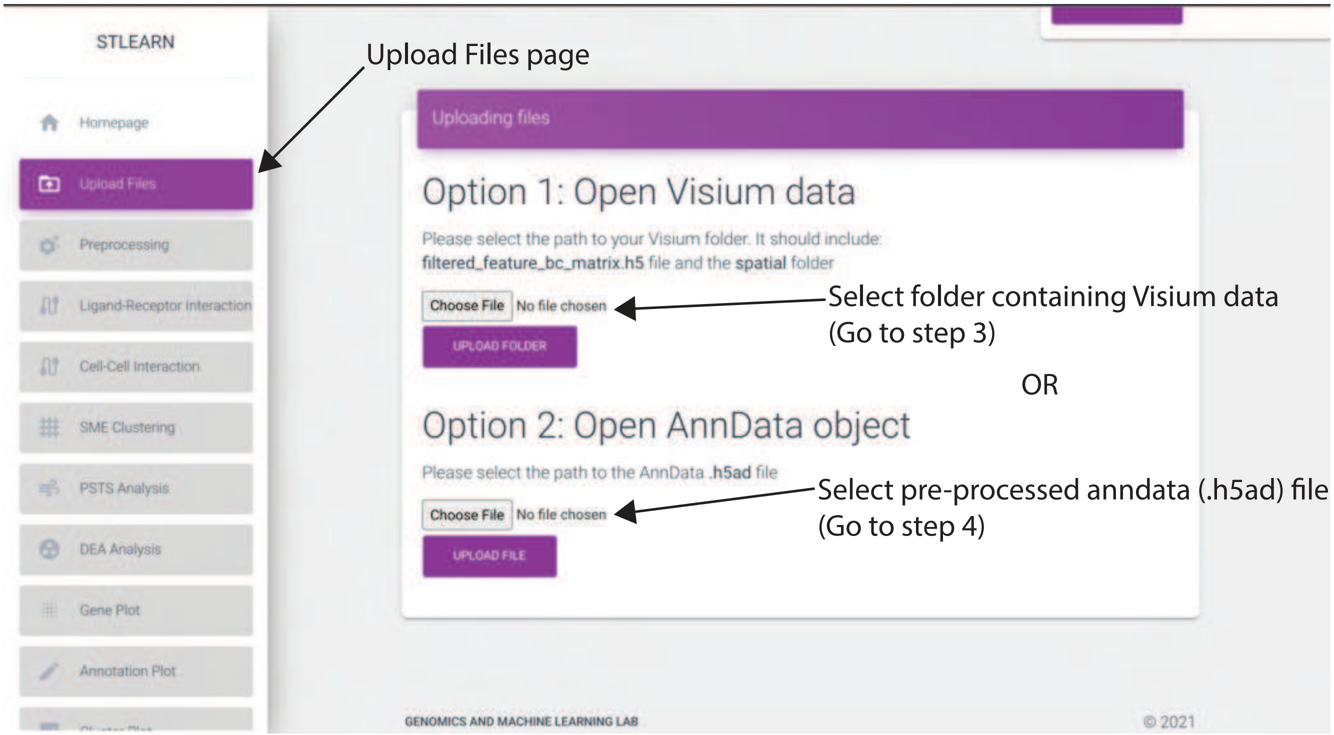


**Supplementary Figure 4.** Step by step instructions for the file uploading function.

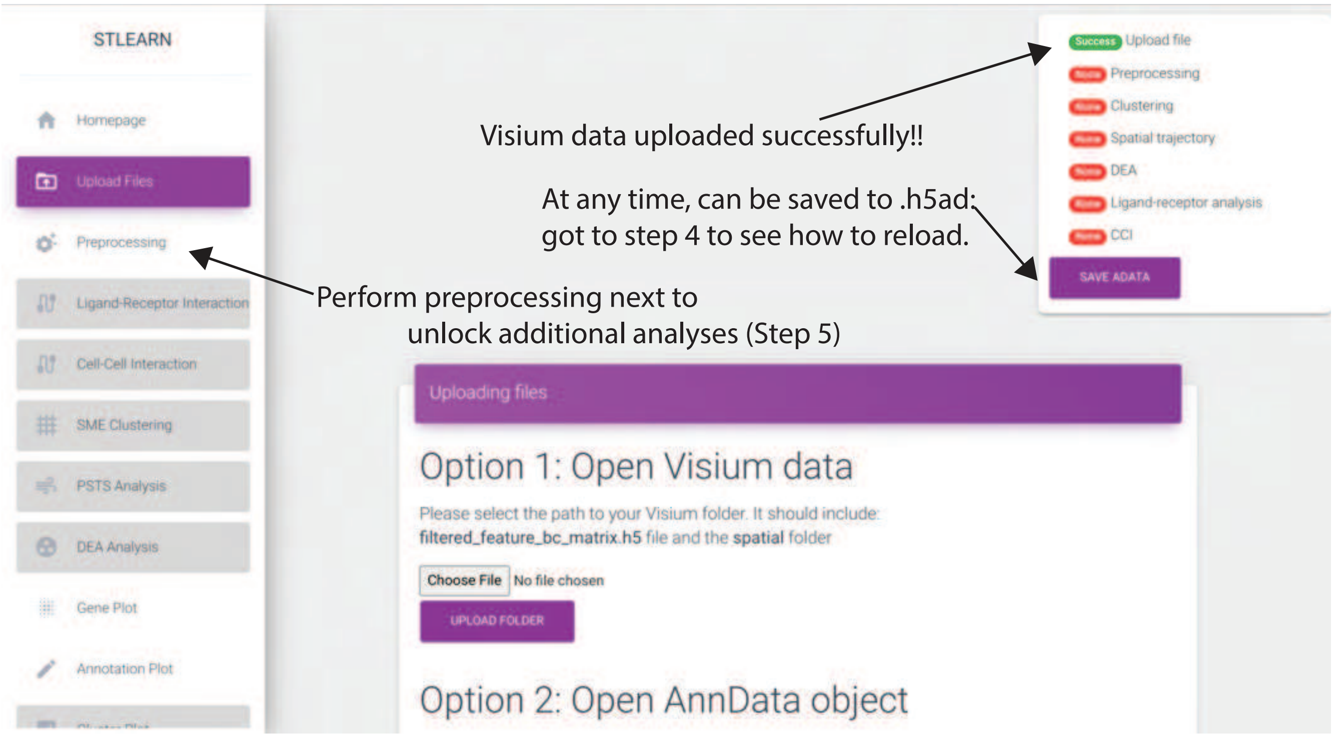


**Supplementary Figure 5.** **Display status afteruploadingthe spatial transcriptomics data files read into AnnData object**.

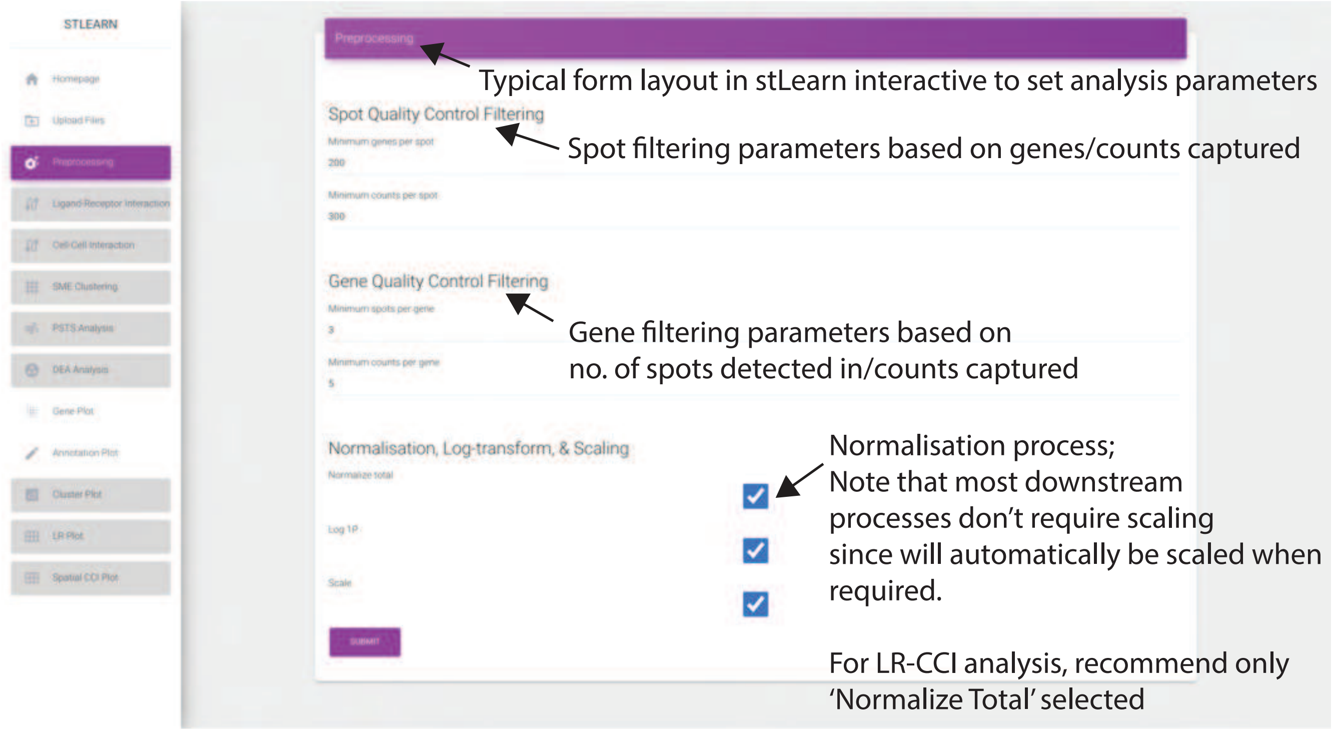


**Supplementary Figure 6.** **Preprocessing options with cell/spot and gene filtering, normalization, log1p transform and data scaling**.

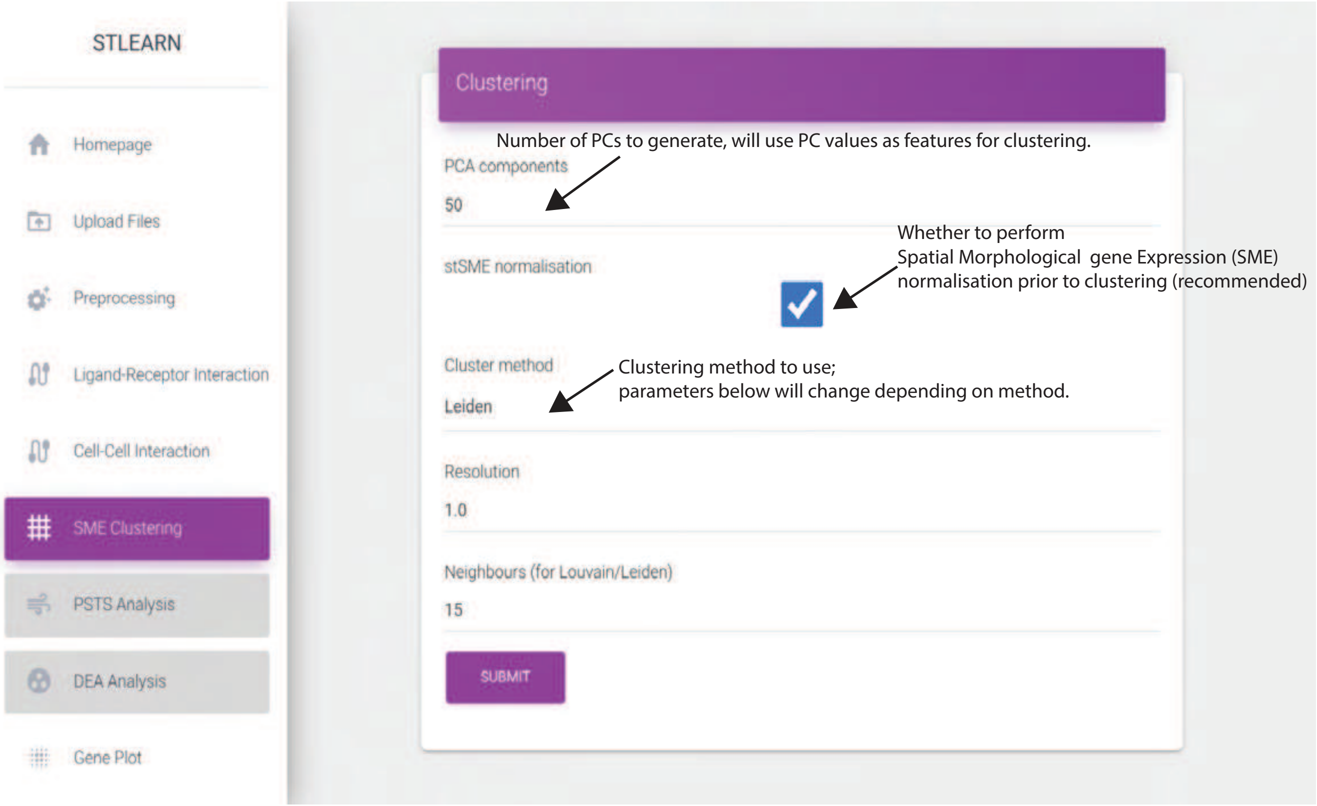


**Supplementary Figure 7.** **Clustering function with PCA setting, SME imputation and clustering method selection.**

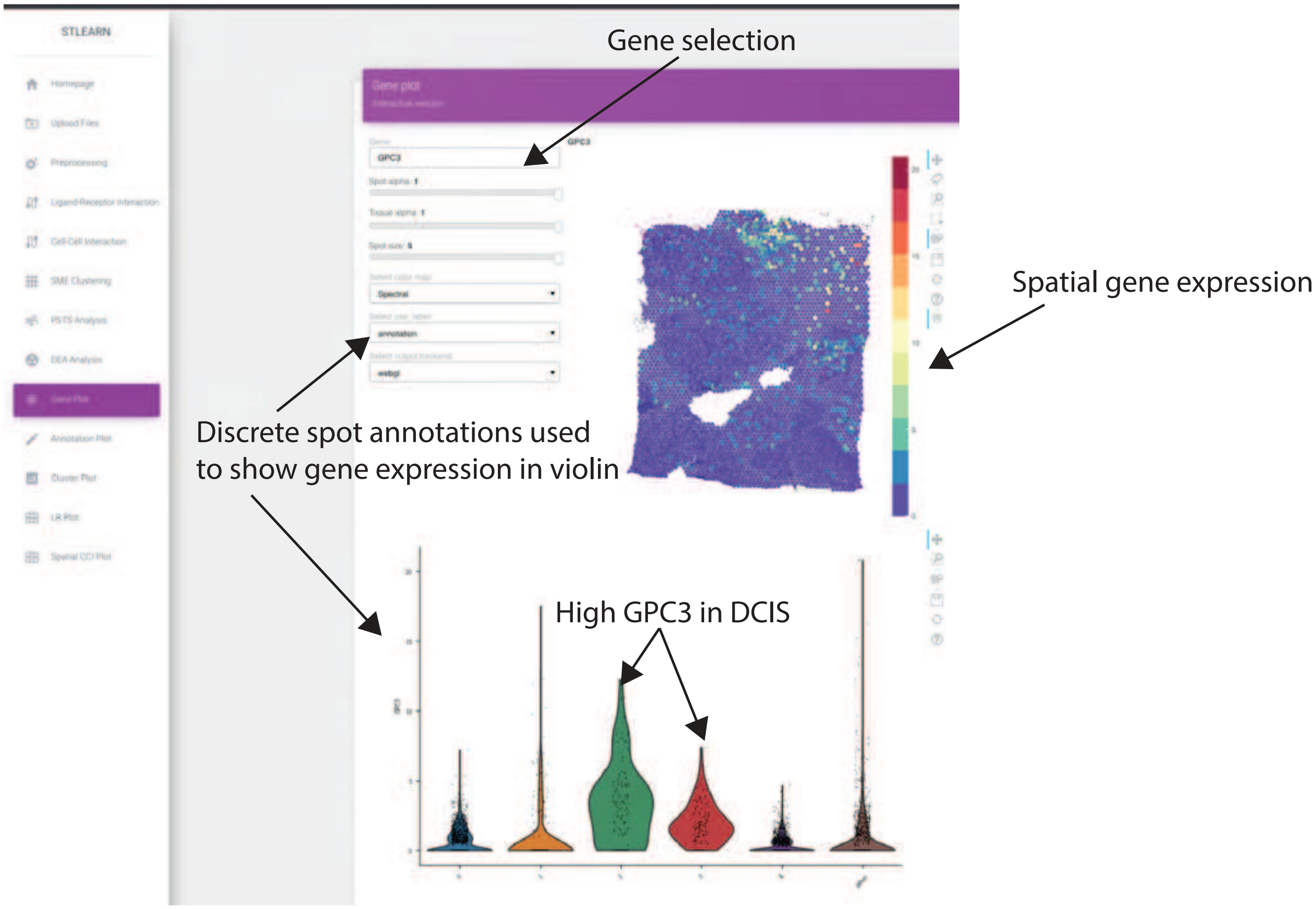


**Supplementary Figure 8.** **Gene expression visualization in the spatial context with different interactive options and tools. i-stLearn also provides a violin plot to show the gene expression distribution across clusters.**

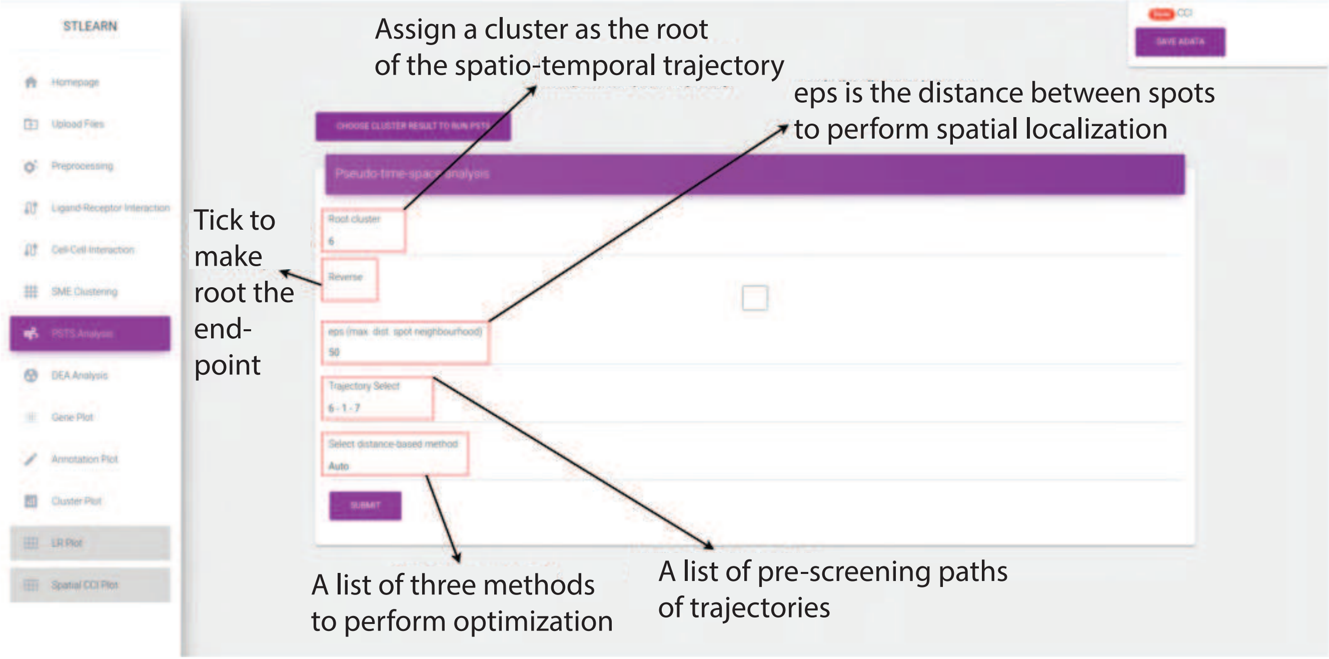


**Supplementary Figure 9.** **Pseudo-time-space analysis function with parameter setting to construct the spatial trajectories**.

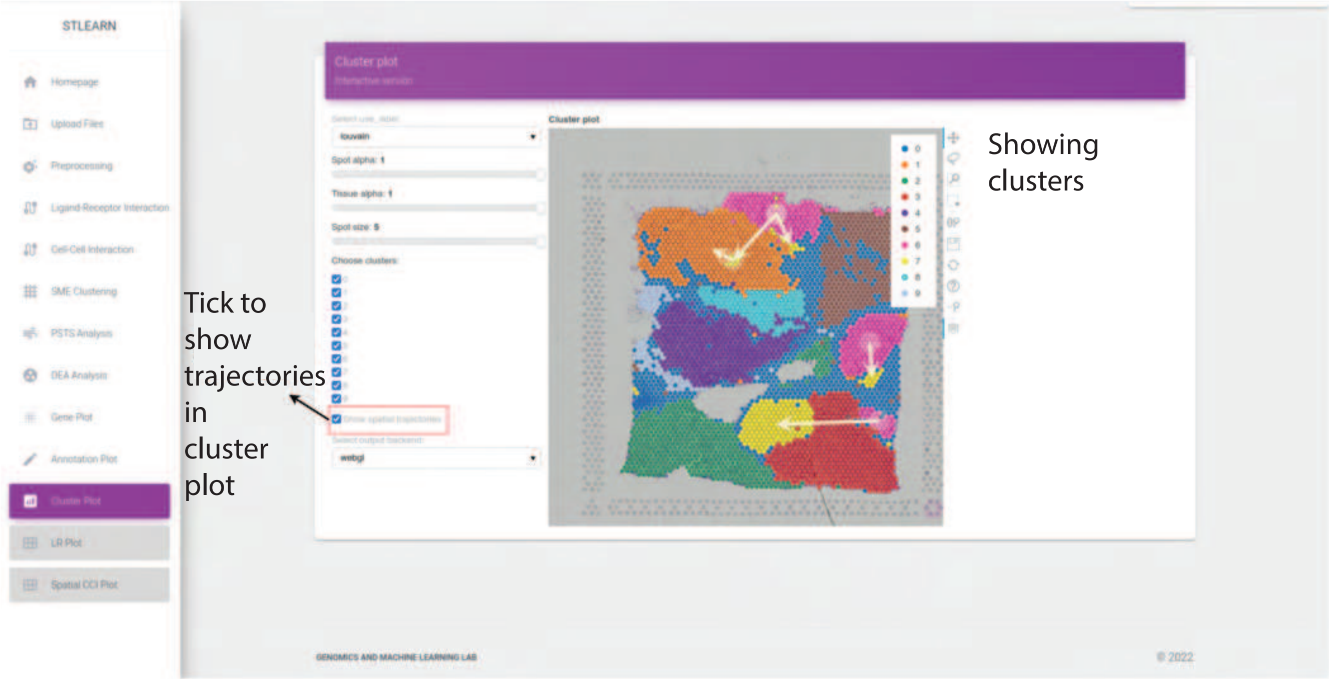


**Supplementary Figure 10.** **Clustering and spatial trajectories visualization of spatial transcriptomics data analysis results**.

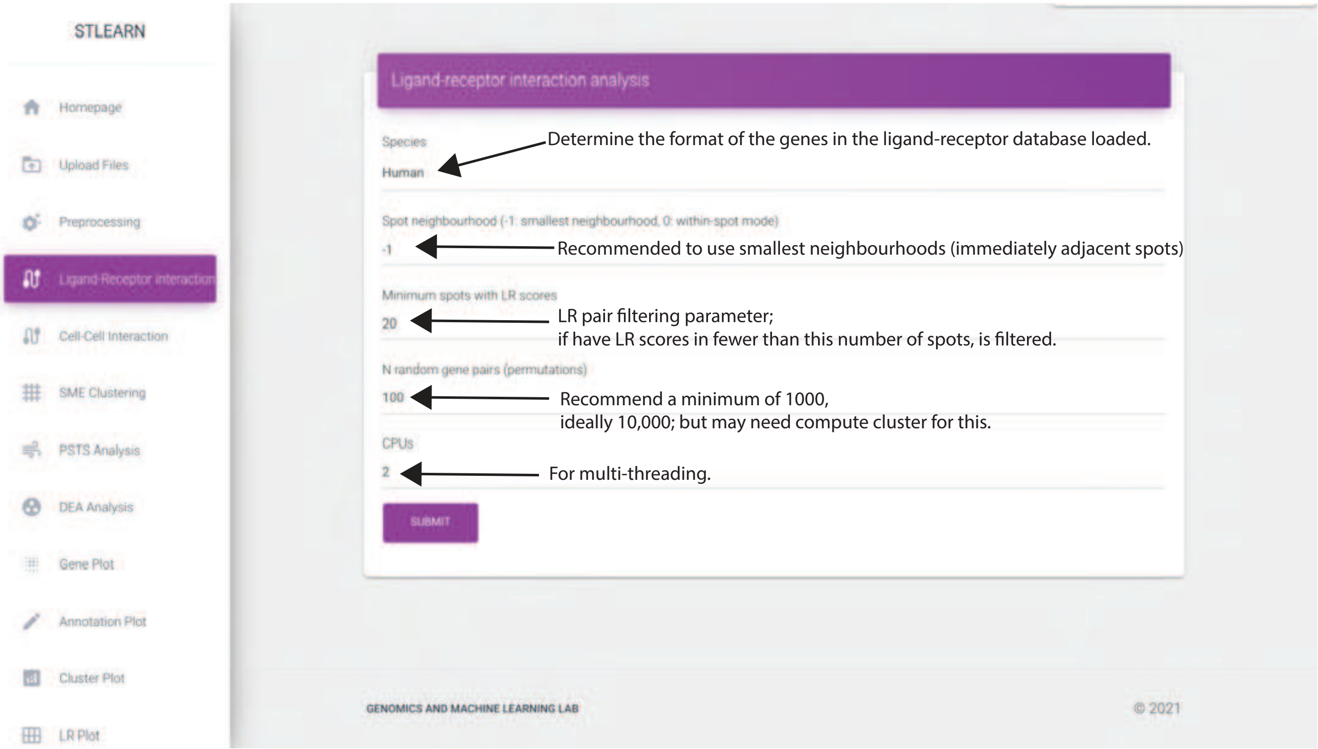


**Supplementary Figure 11.** **Spatial ligand-receptor analysis for cellular communication prediction. Different settings are available**.

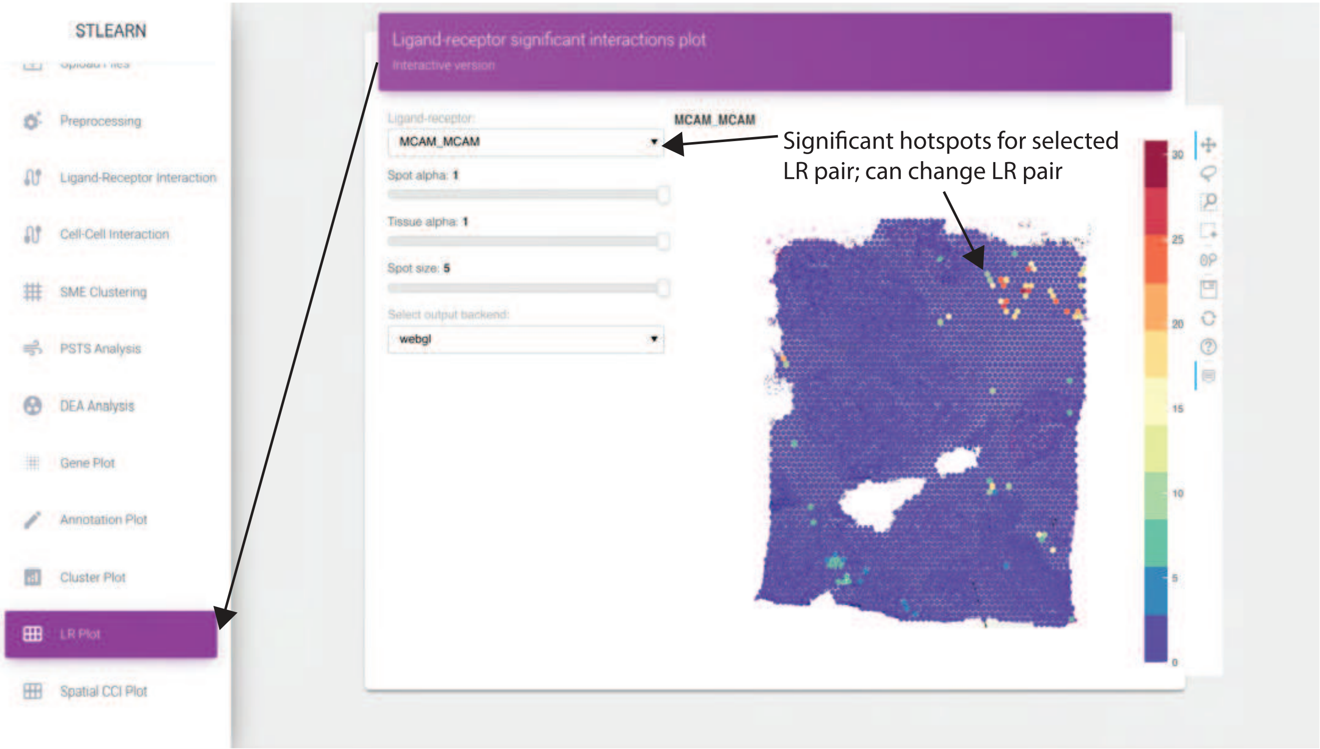


**Supplementary Figure 12. Predicted ligand-receptor interaction results with the LR scores displayed across the breast cancer tissue**.

#### References


1. Sandberg, R. Entering the era of single-cell transcriptomics in biology and medicine. *Nat. methods* **11**, 22–24 (2014).
2. Aldridge, S. & Teichmann, S. A. Single cell transcriptomics comes of age. *Nat. Commun.* **11**, 1–4 (2020).
3. Keren, L. *et al.* A structured tumor-immune microenvironment in triple negative breast cancer revealed by multiplexed ion beam imaging. *Cell* **174**, 1373–1387 (2018).
4. Lewis, S. M. *et al.* Spatial omics and multiplexed imaging to explore cancer biology. *Nat. methods* **18**, 997–1012 (2021).
5. Moses, L. & Pachter, L. Museum of spatial transcriptomics. *Nat. Methods* **19**, 534–546 (2022).
6. Dries, R. *et al.* Advances in spatial transcriptomic data analysis. *Genome research* **31**, 1706–1718 (2021).
7. Pham, D. *et al.* stlearn: integrating spatial location, tissue morphology and gene expression to find cell types, cell-cell interactions and spatial trajectories within undissociated tissues. *BioRxiv* (2020).
8. Wolf, F. A., Angerer, P. & Theis, F. J. Scanpy: large-scale single-cell gene expression data analysis. *Genome biology* **19**, 1–5 (2018).
9. Grinberg, M. *Flask web development: developing web applications with python* (" O'Reilly Media, Inc.", 2018).
10. Jolly, K. *Hands-on data visualization with Bokeh: Interactive web plotting for Python using Bokeh* (Packt Publishing Ltd, 2018).
11. Anders, S. & Huber, W. Differential expression analysis for sequence count data. *Nat. Preced.* 1–1 (2010).
12. Lokhande, P., Aslam, F., Hawa, N., Munir, J. & Gulamgaus, M. Efficient way of web development using python and flask. (2015).
13. Ma, X.-J., Dahiya, S., Richardson, E., Erlander, M. & Sgroi, D. C. Gene expression profiling of the tumor microenvironment during breast cancer progression. *Breast cancer research* **11**, R7 (2009).

**14.** Karaayvaz, M. *et al.* Unravelling subclonal heterogeneity and aggressive disease states in TNBC through single-cell rna-seq. *Nat. communications* **9**, 1–10 (2018).

**15.** Dettogni, R. S. *et al.* Potential biomarkers of ductal carcinoma in situ progression. *BMC cancer* **20**, 119 (2020).

**16.** Traag, V. A., Waltman, L. & Van Eck, N. J. From louvain to leiden: guaranteeing well-connected communities. *Sci. reports* **9**, 1–12 (2019).

**17.** Wolf, F. A. *et al.* PAGA: graph abstraction reconciles clustering with trajectory inference through a topology preserving map of single cells. *Genome biology* **20**, 59 (2019).

**18.** Rodriques, S. G. *et al.* Slide-seq: A scalable technology for measuring genome-wide expression at high spatial resolution. *Science* **363**, 1463–1467 (2019).
